# Discrepancy between *Mtb*-specific IFN-γ and IgG responses in HIV-positive people with low CD4 counts

**DOI:** 10.1101/2022.10.11.511821

**Authors:** Maphe Mthembu, Kathryn A Bowman, Leela RL Davies, Sharon Khuzwayo, Lusanda Mazibuko, Thierry Bassett, Dirhona Ramjit, Zoey Mhlane, Farina Karim, Galit Alter, Thumbi Ndung’u, Emily B Wong

**Author notes:** These authors contributed equally (Thumbi Ndung’u, Emily B Wong).

## Abstract

**Background:** Tuberculosis (TB) is a leading infectious cause of death worldwide and treating latent TB infection (LTBI) with TB preventative therapy is a global priority. This study aimed to measure interferon gamma (IFN-γ) release assay (IGRA) positivity (the current reference standard for LTBI diagnosis) and *Mtb-*specific IgG antibodies in otherwise healthy HIV-negative and HIV-positive adults.

**Methods:** One-hundred and eighteen adults (65 HIV-negative and 53 antiretroviral-naïve HIV-positive), from a peri-urban setting in KwaZulu-Natal, South Africa were enrolled. IFN-γ release following stimulation with ESAT-6/CFP-10 peptides and plasma IgG antibodies specific for multiple *Mtb* antigens were measured using the QuantiFERON-TB Gold Plus (QFT) and customized Luminex assays, respectively. The relationships between QFT status and anti-*Mtb* IgG levels and HIV-status, sex, age and CD4 count were analyzed.

**Findings:** Older age, male sex and higher CD4 count were independently associated with QFT positivity (*p* = 0.045, 0.05 and 0.002 respectively). There was no difference in QFT status between HIV-positive and HIV-negative groups (58% and 65% respectively, *p* = 0.06), but within CD4 count quartiles, people with HIV had higher QFT positivity than people without HIV (*p* = 0.008 (2^nd^ quartile), <0.0001 (3^rd^ quartile)). *Mtb*-specific IFN-γ levels were lowest, and *Mtb*-specific IgGs were highest in HIV-positive individuals with the lowest CD4 counts.

**Interpretation:** These results suggest that the QFT assay underestimates LTBI among immunosuppressed people with HIV and *Mtb*-specific IgG may be a useful alternative biomarker for *Mtb* infection. Further evaluation of how *Mtb*-specific antibodies can be leveraged to improve LTBI diagnosis is warranted, particularly in HIV-endemic areas.

**Funding:** The study was funded by the NIH/NIAID [K08AI118538] (EBW) and, in part, by the Africa Health Research Institute through the Wellcome [Strategic Core award: 201433/Z/16/A]. The study was also supported in part by the Strategic Health Innovation Partnerships (SHIP) Unit of the South African Medical Research Council with funds received from the South African Department of Science and Innovation as part of a bilateral research collaboration agreement with the Government of India. Other support came from the South African Research Chairs Initiative and the Victor Daitz Foundation (TN) and the Burroughs Wellcome Fund Investigators in Pathogenesis of Infectious Disease [1022002] (EBW). This research was also funded in part by the Sub-Saharan African Network for TB/HIV Research Excellence (SANTHE) through a grant [DEL-15-006] by the Wellcome Trust and the UK Foreign, Commonwealth & Development Office, with support from the Developing Excellence in Leadership, Training and Science in Africa (DELTAS Africa) programme (TN, MM). For the purpose of open access, the author has applied a CC BY public copyright license to any Author Accepted Manuscript version arising from this submission.

**Copyrights:** © 2022 The Authors. Published by Elsevier B.V. This is an open access article under the CC BY-NC-ND license (https://creativecommons.org/licenses/by-nc-nd/4.0/)

**Research in context:** *Evidence before this study:* *Mtb*-specific IFN-γ production as measured by IGRA is the current gold standard for determining latent TB infection. However, since these tests measure immunoreactivity to Mtb peptides, they are indirect measures of *Mtb* infection and their performance characteristics are impacted by co-infections and comorbidities that influence immune responses, including HIV. Recently, a human phenotype has been defined in people who are highly exposed to *Mtb* but consistently test negative for evidence of *Mtb* infection by IGRA and tuberculin skin test (TST). These individuals have been observed to have a unique profile of *Mtb*-specific antibodies when compared to the classical IGRA positive LTBI group, suggesting that *Mtb*-specific antibodies may identify additional people with a history of *Mtb* infection or exposure when compared to IGRA alone. Comparison of IGRA and Mtb-specific antibodies in people living with HIV has not previously been performed.

*Added value of this study:* Here, we concurrently assessed *Mtb*-specific IFN-γ production and IgG in a cohort of 118 well-defined HIV-negative and antiretroviral naïve HIV-positive individuals from KwaZulu-Natal, South Africa, a highly TB endemic area. We found a discrepancy between *Mtb*-specific IFN-γ and *Mtb*-specific IgG levels, particularly in HIV-positive individuals with low CD4 cell counts. Notably people with the lowest CD4 counts had the highest levels of *Mtb*-specific IgG levels in the plasma, and the lowest levels of QTF positivity.

*Implications of all evidence available:* IGRAs may underestimate *Mtb* infection status, especially in people with HIV infection or who have T cell depletion or dysfunction. *Mtb*-specific IgG antibodies indicate development of a B cell response to *Mtb* and may have promise as an alternative biomarker of TB immunoreactivity that does not depend on T cell function.

## Introduction

Tuberculosis (TB) is the leading cause of death for people living with HIV (PLWH). Despite recent improvements in TB diagnostics and increasing access to anti-tubercular therapy, TB mortality remains high with an estimated annual death rate of 15% of the 10 million annual TB incidents, with 21% of those deaths attributable to co-infection with HIV (1). An important challenge to eliminating TB is that *Mycobacterium tuberculosis* (*Mtb*) infection presents as a dynamic continuum, ranging from rapidly cleared *Mtb* infection to active TB (ATB) disease. Currently available diagnostic tools are imperfect and are incapable of accurately identifying the different stages of TB pathogenesis (2, 3). Considering that there are no direct tests of mycobacterial burden, people who have immunological evidence of *Mtb* infection and who are clinically free of disease have historically been considered to have “latent TB infection” (LTBI), although use of this term is increasingly criticized because of its inherent imprecision (4–6). It has been estimated that 30% of people in the world have LTBI and PLWH who have LTBI are at higher risk of progressing to active disease (1, 7). Preventative therapies, including 6-9 months of isoniazid and more recently tested shorter courses of multiple drugs, like 3 months of rifapentine and isoniazid (3HP), have been successful in reducing the short-term risk of progression to ATB in PLWH (8–10).

The impact of TB prevention therapies has been limited by the indirect nature of diagnostic tests for LTBI. The tuberculin skin test (TST) uses purified protein derivatives (PPD) from *Mtb* and has significant rates of cross-reactivity with the *M. bovis* BCG strain used to vaccinate infants in TB endemic countries (5). *Mtb*-specific IGRA tests have improved specificity for *Mtb* and are therefore the current gold standard for determining *Mtb* infection status, however, both tests (TST and IGRA) rely on T cell immune responses (4, 5, 11) and therefore fail to determine whether viable *Mtb* infection is present. Additionally, both tests perform less well in immunocompromised people with decreased T cell function, including PLWH (12, 13). Moreover, the indirect nature of these tests limits the field’s ability to precisely define the pathogenesis of LTBI (14), which is required to design more effective TB prevention strategies that are targeted to the optimal stage in the spectrum of TB disease.

In this study, we sought to measure rates of and characteristics associated with LTBI in a peri-urban setting in KwaZulu-Natal, South Africa by comparing *Mtb*-specific IFN-γ and *Mtb*-specific IgG antibodies in otherwise healthy HIV-negative and HIV-positive adults.

## Methods

### Study population and clinical characteristics

Data and plasma from participants who were screened for the Phefumula cohort, a research bronchoscopy study described previously in Muema *et al*, 2020 (15) and Khuzwayo *et al*, 2021 (16), were used for this analysis. HIV-positive and HIV-negative adults, between 18-50 years, who were otherwise healthy (no chronic medical conditions, no current or past tobacco use, no current TB symptoms) were recruited from KwaDabeka Community Health Clinic in peri-urban KwaZulu-Natal, South Africa. HIV-positive participants were newly diagnosed and were antiretroviral therapy naïve. The HIV status of all participants was determined by fourth generation enzyme linked-immunosorbent assay (ELISA) testing and HIV RNA plasma viral load (Sigma Aldrich).

### Ethics Statement

All participants provided written informed consent for participation. Study protocols were approved by the University of KwaZulu-Natal Biomedical Research Ethics Committee (BREC, protocol BF503/15), the Partners Institutional Review Board and the University of Alabama Institutional Review Board.

### Measurement of *Mtb*-specific IFN-γ

The QuantiFERON-TB Gold Plus (QFT-Plus) assay (Qiagen) was conducted as per manufacturer’s instructions. Briefly, whole blood was collected into QFT tubes at room temperature, thoroughly mixed and then incubated at 37°C for 20 hours to allow stimulation by TB antigens (early secretory antigenic target 6 [ESAT6] and culture filtrate protein 10 [CFP10]). After centrifugation at 2,500 RPM for 15 minutes, supernatant was collected and 50 μL used for IFN-γ ELISA, performed according to the manufacturer’s instructions. The results were analysed using the QFT-Plus analysis software v2.71.2 (Qiagen). A positive QuantiFERON was defined by IFN-γ produced in response to TB1 or TB2 minus the NIL tube (negative control, ≤ 0.8 IU/mL) ≥ 0.35 IU/mL.

### Measurement of *Mtb*-specific IgG

#### Antigen selection and source

Purified protein derivative (PPD) was obtained from Staten Seruminstitut. Antigen 85 complex (Ag85), ESAT6, CFP10, heat shock protein X (HspX), alanine- and proline-rich secreted protein (Apa), groES, crystallin, and lipoarabinomannan (LAM) were obtained from BEI Resources (NR-14855, NR-49424, NR-49425, NR-49428, NR-14862, NR-14861, NR-14860, NR-14848, respectively). Influenza hemagglutinin from A/New Caledonia/20/1999 and B/Brisbane/60/2008 (ImmuneTech) was included in all Luminex assays as an internal control.

#### Customized multiplex Luminex assay

*Mtb*-specific IgG levels were measured using a Luminex isotype assay described by Lu *et al*, 2019 (17). Briefly, we bound Luminex MagPlex® carboxylated beads (Luminex) with *Mtb*-specific antigens and non-*Mtb* related antigen using NHS-ester linkages following manufacturer’s recommendations. Protein antigens were coupled to Magplex carboxylated beads using 1-ethyl-3-(3-dimethylaminopropyl) carbodiimide hydrochloride and N-hydroxysulfosuccinimide) according to the manufacturer’s recommendations. LAM was modified by 4-(4,6-dimethoxy[1,3,5]triazin-2-yl)-4-methyl-morpholinium prior to conjugation. Multiplexed antigen-coupled beads were incubated with sample serum at a 1:100 dilution in assay buffer (PBS with 0.1% BSA and 0.05% Tween) for 18 hours. IgG was detected using phycoerythrin-conjugated mouse antibodies against human antibody subclasses (Southern Biotech). Secondary incubations were performed over 2 hours. Flow cytometric analysis was performed on an iQue Screener PLUS using ForeCyt software (Intellicyt). IgG level concentration was expressed as Median Fluorescence Intensity (MFI) and absolute concentrations of these was not calculated. All assays were performed in duplicates.

### Quantitative measurement of total IgG

ELISA plates were coated overnight with anti-human IgG (Sigma Aldrich I5260) at 1:5 000 in PBS, then washed 3 times in PBS-0.05% Tween (PBST) and blocked with 5% BSA in PBS. Primary incubation was performed with serum diluted at 1:1 000 000 for 2 hours at room temperature, then plates were washed 3 times in PBST. Secondary incubation was performed with 1:20 000 dilution of HRP-conjugated anti-human IgG antibody (Sigma Aldrich SAB701283) for 1 hour at room temperature, then plates washed 3 times in PBST. Plates were developed with 1-Step™ Ultra TBM-ELISA substrate solution (Thermo Fisher) and stopped with equal volume 1N H_2_SO_4_. Absorbance was read at 450 nm with reference 570 nm. GammaGard liquid IVIG was used as a quantitative standard.

### Statistical analyses

Statistical analyses and graphical representations were performed using GraphPad Prism v9.2.0 (GraphPad Software) and Stata/SE v17.0 (StataCorp LLC). Non-parametric tests (Fishers exact and Chi-squared tests for single or multiple categorical comparisons and Mann-Whitney *U* and Kruskal-Wallis tests for single and multiple comparisons of continuous dependent variables.) were used to avoid assumptions about data normality. Logistic regression analysis was employed to assess multivariate analysis of QFT status as a binary dependent variable. For single comparisons a significance threshold of *p* < 0.05 was used. For measures of correlations between study variables, Benjamini-Hochberg correction tests were applied to account for multiple comparisons.

## Results

### Characteristics of study participants

One hundred and eighteen otherwise healthy adults between ages of 18-50 years were enrolled. Sixty-five of them were HIV-negative, and 53 were HIV-positive (Table 1). To meet enrollment criteria, participants had to be feeling well, have no prior history of TB or other co-morbidity. All the HIV-positive participants were newly diagnosed and antiretroviral therapy (ART)-naïve. The majority of participants were female (69%), with sex distribution balanced between the HIV-positive (71% females) and HIV-negative groups (66% females) (*p* = 0.6969). The HIV-positive group was slightly older than the HIV-negative group (*p* = 0.0234). CD4 T-cell count was measured for all study participants and the median (interquartile range, IQR) for the HIV-positive and HIV-negative participants was 358 cells/mm^3^ (IQR: 210 - 575) and 870 cells/mm^3^ (IQR: 710 - 1 201), respectively (*p* < 0.0001). Viral load for the HIV-positive group was high (median of 123 919 copies/mL (IQR: 10 600 - 2 124 982)), as expected for an ART-naïve population. Age and CD4 count values of all participants were stratified into quartiles and the number of participants in each quartile for the HIV-positive and HIV-negative groups is shown in Table 1.

**Table 1:**
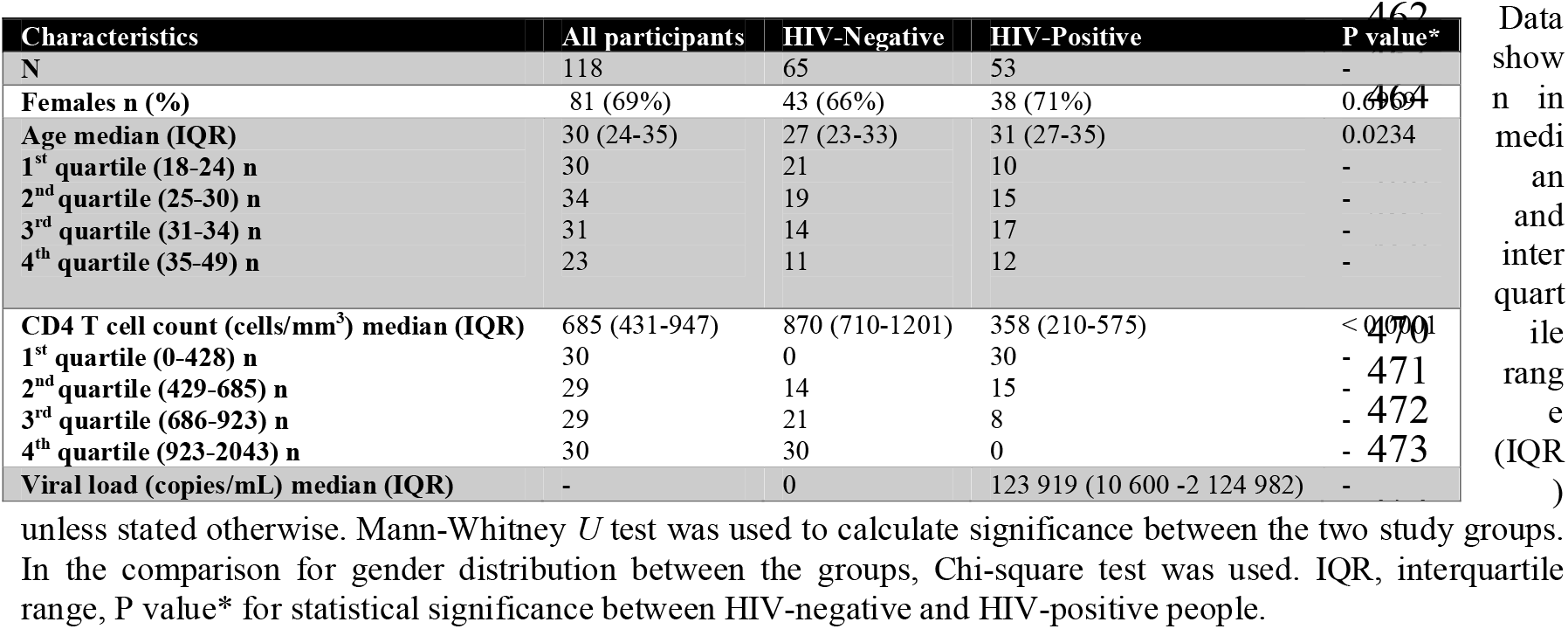
Demographic and clinical characteristics.

### ESAT-6/CFP-10-specific IFN-γ is associated with higher CD4 cell count

We first aimed to determine the rates of QFT positivity within our study population. All study participants were screened for latent TB infection using the QFT-Gold plus test. Fifty-eight percent of participants had a positive QFT test with a trend towards higher positivity among the HIV-negative (65%) compared to the HIV-positive (51%) group (*p* = 0.06) (Figure 1A). Stratifying participants by age quartiles showed a significant increase in QFT positivity with increasing age (*p* = 0.006) (Figure 1B). Males had higher QFT positivity compared to females (*p* = 0.0419) (Figure 1C). Stratification by CD4 cell count quartiles showed that QFT positivity increased significantly with CD4 cell count (*p* < 0.0001) (Figure 1D).

**Figure 1:**
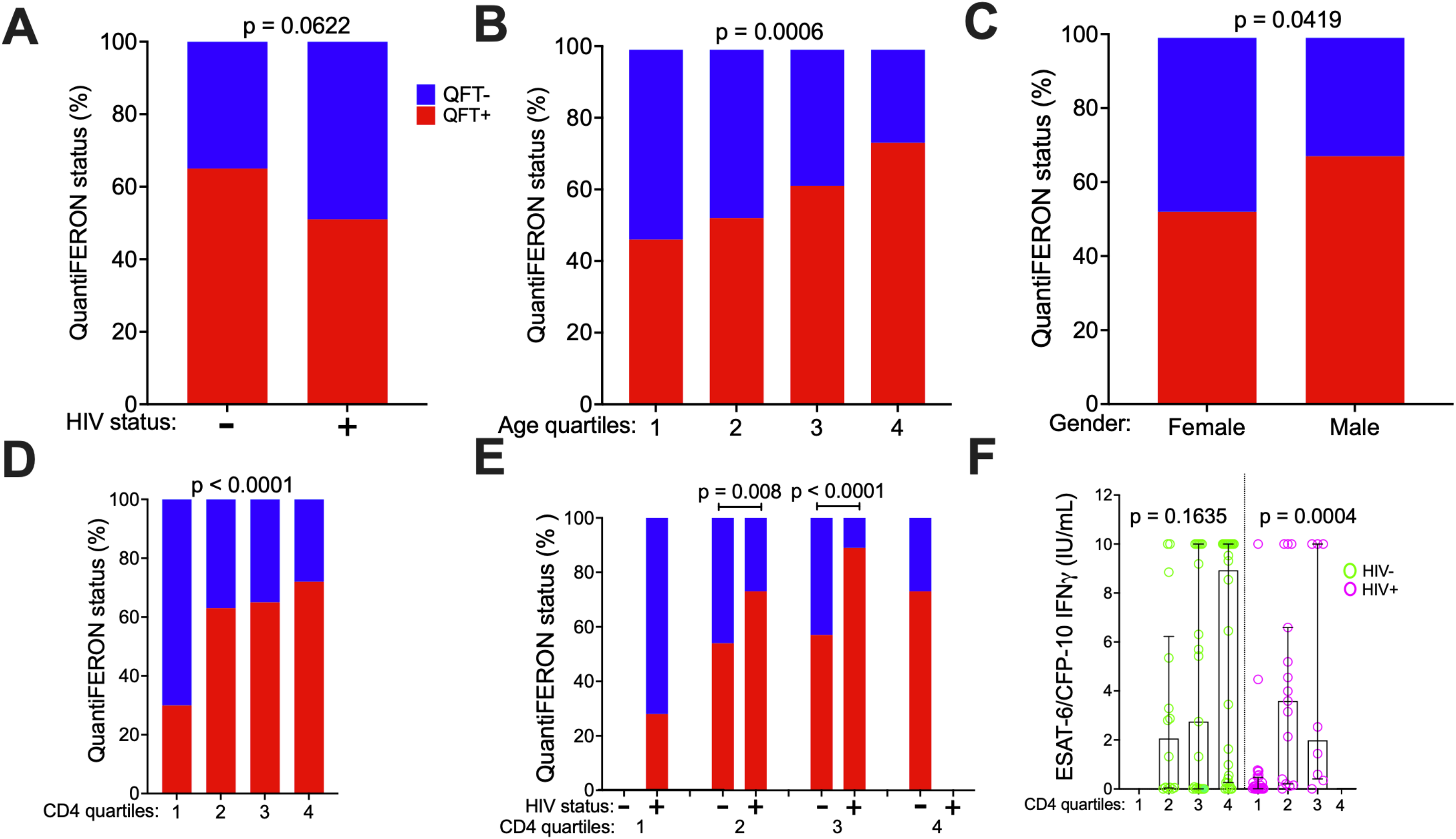
Factors associated with QuantiFERON positivity. **A** Participants (n = 118) stratified according to HIV status and QuantiFERON test results expressed as percentage (%) positive (red bars) or negative (blue bars) per group. Fisher's Exact test was used to test for significant difference between the 2 groups **B** Participants’ age stratified into quartiles (1 = 18 - 24: 2 = 24 – 30: 3 = 30 – 35: 4 = 35 – 39) and QuantiFERON test results expressed as percentage positive (red bars) or negative (blue bars) per age quartile. Chi squared test was used to test for significant differences across all quartiles. **C** The participants were also stratified by sex and QuantiFERON status. Fisher's Exact test was used to test for significant difference between males and females. **D** The CD4 count was stratified into quartiles (1 = 0 - 428: 2 = 429 - 685: 3 = 686 - 923: 4 = 924 - 2043 all in cells/mm^3^ and QuantiFERON status expressed as percentage positive (red bars) or negative (blue bars) per quartile. The Chi-squared test was used to test for significant difference across all CD4 count quartiles. **E** QuantiFERON status for each CD4 count quartile in HIV− (n = 65) versus HIV+ (n = 53) participants. Fishers Exact test was used to test for significant differences between comparable CD4 count quartiles, 2 and 3. **F** ESAT-6/CFP-10 specific IFN-γ production among participants in the different CD4 counts quartiles, stratified by HIV− (green) and HIV+ (pink).

We hypothesized that people with HIV would have equivalent or higher rates of latent TB infection compared to HIV-negative people, and that CD4 T-cell deficiency led to the lower rates of positive QFT testing observed in HIV-positive individuals. To test this hypothesis, we stratified the entire cohort into quartiles by CD4 T-cell count. Quartile 1 included only HIV-positive individuals and quartile 4 included only HIV-negative individuals; Quartiles 2 and 3 each contained HIV-positive and HIV-negative individuals. When compared within CD4 quartile, HIV-positive people had higher QFT positivity than HIV-negative people (*p* = 0.008, and *p* < 0.0001 for quartiles 2 and 3 respectively, Figure 1E). Among HIV-negative participants, there were no differences in the production of IFN-γ across CD4 count quartiles (*p* = 0.1635, Figure 1F). However, in the HIV-positive group there was significantly lower production of IFN-γ in the lowest CD4 count quartile (*p* = 0.0004, Figure 1F). In a multivariate analysis that accounted for age, sex, HIV status and CD4 count, older age (*p* = 0.045), male sex (*p* = 0.05) and higher CD4 cell count (*p* = 0.002) were independently associated with QFT positivity and HIV-status was not (Table 2). Taken together, these data indicate that CD4 count is associated with lower quantitative IFN-γ release in HIV-positive individuals, and may underlie the lower rates of QFT positivity observed in this population.

**Table 2:**
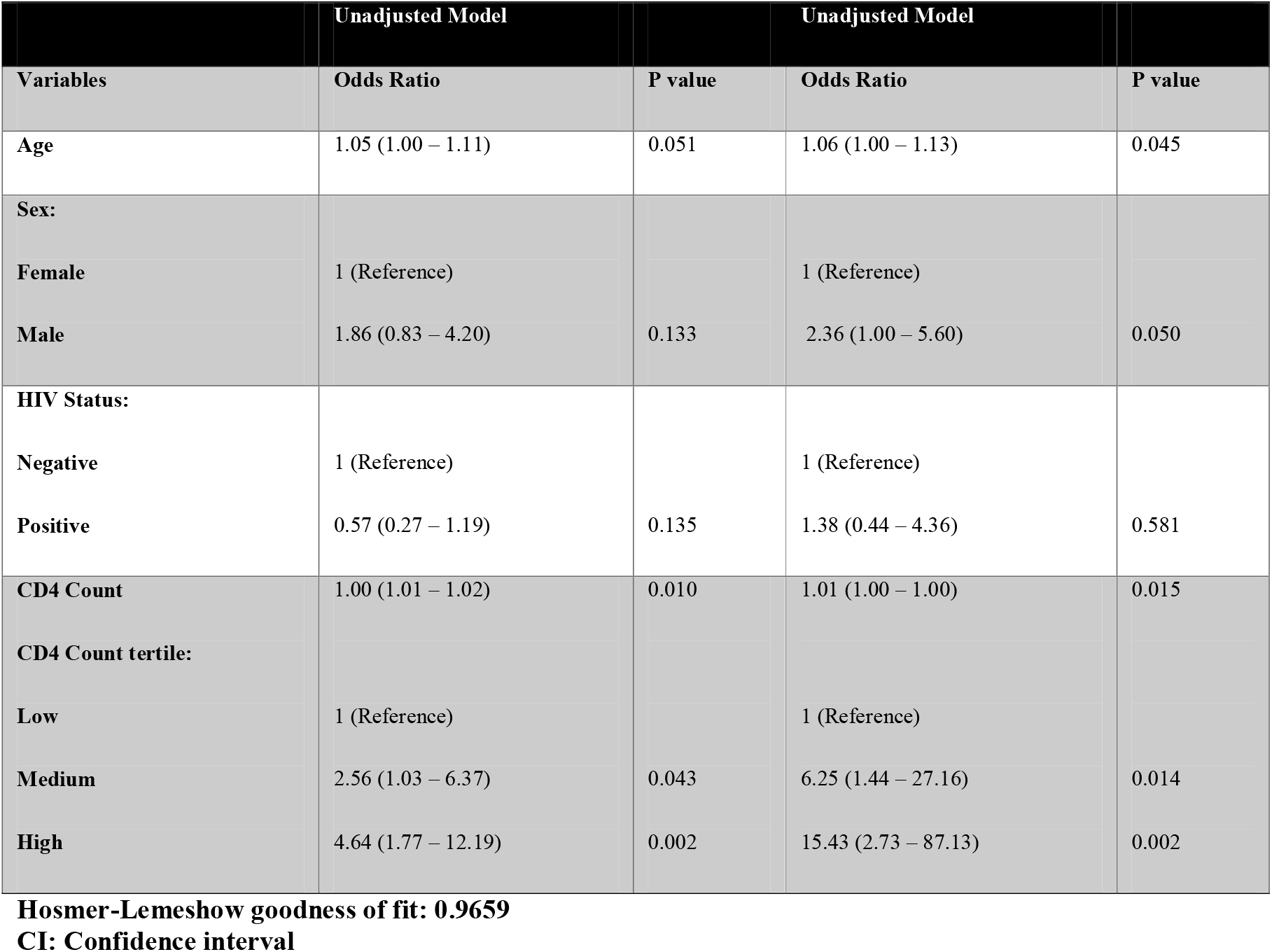
Unadjusted and adjusted multivariate logistic regression model for QuantiFERON positivity.

| Unadjusted Model |  |  | Unadjusted Model |  |
| --- | --- | --- | --- | --- |
| Variables | Odds Ratio | P value | Odds Ratio | P value |
| Age | 1.05 (1.00 – 1.11) | 0.051 | 1.06 (1.00 – 1.13) | 0.045 |
| Sex: |  |  |  |  |
| Female | 1 (Reference) |  | 1 (Reference) |  |
| Male | 1.86 (0.83 – 4.20) | 0.133 | 2.36 (1.00 – 5.60) | 0.050 |
| HIV Status: |  |  |  |  |
| Negative | 1 (Reference) |  | 1 (Reference) |  |
| Positive | 0.57 (0.27 – 1.19) | 0.135 | 1.38 (0.44 – 4.36) | 0.581 |
| CD4 Count | 1.00 (1.01 – 1.02) | 0.010 | 1.01 (1.00 – 1.00) | 0.015 |
| CD4 Count tertile: |  |  |  |  |
| Low | 1 (Reference) |  | 1 (Reference) |  |
| Medium | 2.56 (1.03 – 6.37) | 0.043 | 6.25 (1.44 – 27.16) | 0.014 |
| High | 4.64 (1.77 – 12.19) | 0.002 | 15.43 (2.73 – 87.13) | 0.002 |

Hosmer-Lemeshow goodness of fit: 0.9659 CI: Confidence interval

### ESAT-6/CFP-10-specific antibody responses are higher in HIV-positive people with low CD4 cell counts

Since IGRAs rely on functional CD4 T-cell responses, we next asked whether antibody responses could provide a more sensitive marker of *Mtb* exposure or infection in HIV-positive individuals. We measured the *Mtb*-specific IgG responses in the same individuals using a customised Luminex assay to characterize the *Mtb*-specific antibody response (17). ESAT-6/CFP-10-specific IgG was analysed according to HIV status, age, sex and CD4 count. ESAT-6/CFP-10-specific IgG levels were significantly higher in HIV-positive people (*p* < 0.0001) compared to HIV-negative (Figure 2A). In contrast, levels of influenza hemagglutinin (HA)-specific IgG (non-*Mtb* control) did not differ by HIV status (*p* = 0.5883) (Figure 2A). Levels of ESAT-6/CFP-10-specific IgG increased with age (*p* = 0.0256) (Figure 2B) but did not differ between males and females (*p* = 0.3286) (Figure 2C).

**Figure 2:**
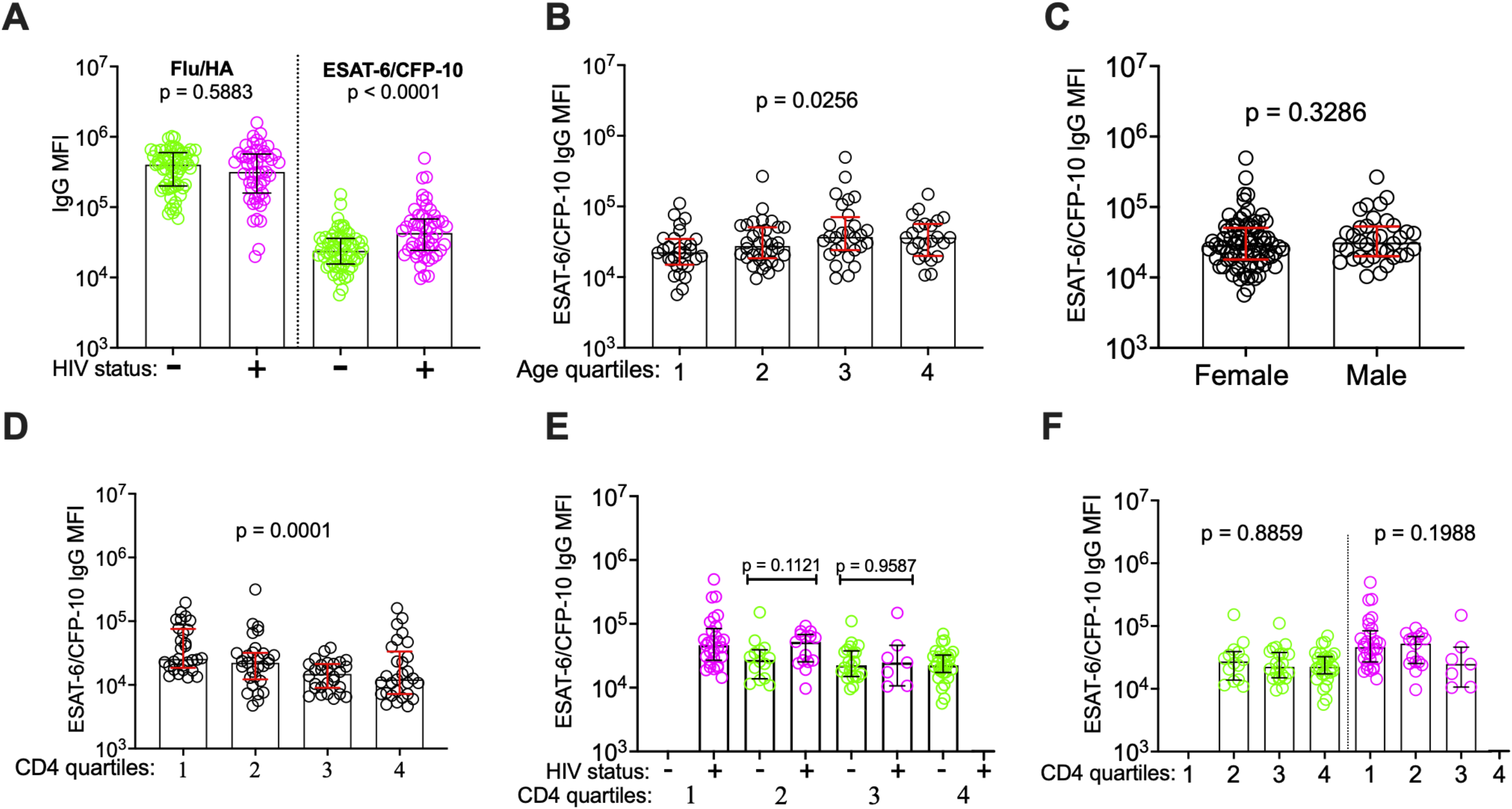
ESAT-6/CFP-10 specific total IgG and how it associates with QuantiFERON status associated factors. **A.** Influenza HA (non-*Mtb* related antigen) and ESAT-6/CFP-10 total IgG expression stratified by HIV status. Mann-Whitney *U* used to test for statistical significance between HIV− (green circles) and HIV+ (pink circles) groups. **B.** Expression of ESAT-6/CFP-10 total IgG across age quartiles (1 = 18 - 24: 2 = 24 – 30: 3 = 30 – 35: 4 = 35 – 39) of all study participants. Kruskal-Wallis used to test for statistical significance across age quartiles. **C.** Expression of ESAT-6/CFP-10 total IgG by sex. Mann-Whitney *U* used to test for statistical significance between the groups. **D.** Expression of ESAT-6/CFP-10 total IgG across all participants stratified by CD4 count quartiles. Kruskal-Wallis test used to test for statistical significance across quartiles. **E.** ESAT-6/CFP-10 total IgG across CD4 count quartiles (1 = 0 - 428: 2 = 429 - 685: 3 = 686 - 923: 4 = 924 - 2043 all in cells/μ L) by HIV status (green: HIV− and pink: HIV+) and Mann-Whitney *U* was used to test for statistical significance across quartiles. **F.** Expression of ESAT-6/CFP-10 total IgG across all participant CD4 T cell count quartiles stratified by HIV status (green: HIV− and pink: HIV+). (n=115, HIV− n= 64, HIV+ n= 51).

We next asked whether *Mtb*-specific IgG responses differed across CD4 T-cell count quartiles. ESAT-6/CFP-10-specific IgG levels differed significantly by CD4 count quartiles (*p* = 0.0001), with participants in the lowest CD4 count quartile having the highest IgG levels (Figure 2D). In contrast to the pattern observed for ESAT-6/CFP-10 IFN-γ, HIV status did not significantly affect the expression of ESAT-6/CFP-10-specific IgG when HIV-negative and HIV-positive participants in the 2^nd^ and 3^rd^ CD4 cell count quartiles were compared (*p* = 0.1121 and *p* = 0.9587 respectively) (Figure 2E). Within both the HIV-negative and the HIV-positive sub-populations there was no significant difference in ESAT-6/CFP-10-specific IgG between CD4 count quartiles (*p* = 0.8859 and *p* = 0.1988, Figure 2F). Thus, while ESAT6/CFP10-specific IgG is higher in HIV-positive individuals in our cohort, this association is unrelated to quantitative CD4 T-cell count.

### Additional anti-*Mtb* specific IgG antibody responses are also highest in HIV-positive people with low CD4 cell counts

To understand whether the observed patterns for ESAT-6/CFP-10-specific IgG levels were consistent for other anti-*Mtb* IgG antibodies, we next measured levels of antibodies to 7 other well-characterized and immunogenic *Mtb* antigens (Apa, HspX, GroES, alpha-crystallin, Ag85a/b, PPD, LAM) and analysed them by HIV-status and CD4 cell count quartile. Six of the 7 measured *Mtb*-specific IgG responses were significantly increased in HIV-positive people, including Apa-specific IgG (*p* = 0.0003), GroES-specific IgG (*p* < 0.0001), HspX-specific IgG (*p* < 0.0001), Crystallin-specific IgG (*p* = 0.0009), Ag85a/b-specific IgG (*p* < 0.0001), anti-PPD IgG (*p* < 0.0001). LAM-specific IgG was the only *Mtb*-specific IgG that did not differ by HIV status (*p* = 0.1991) (Figure 3B-H). Similarly, 6 of the 7 measured *Mtb*-specific antigens significantly differed across CD4 count quartile (Apa IgG-specific (*p* = 0.0001), GroES-specific IgG (*p* = 0.0011), HspX IgG-specific (*p* < 0.0001), Crystallin-specific IgG (*p* = 0.0457), Ag85a/b-specific IgG (*p* = 0.0001), PPD-specific IgG (*p* < 0.0001)), with a LAM-specific IgG again the exception (*p* = 0.7146, Figure 3I-P).

**Figure 3:**
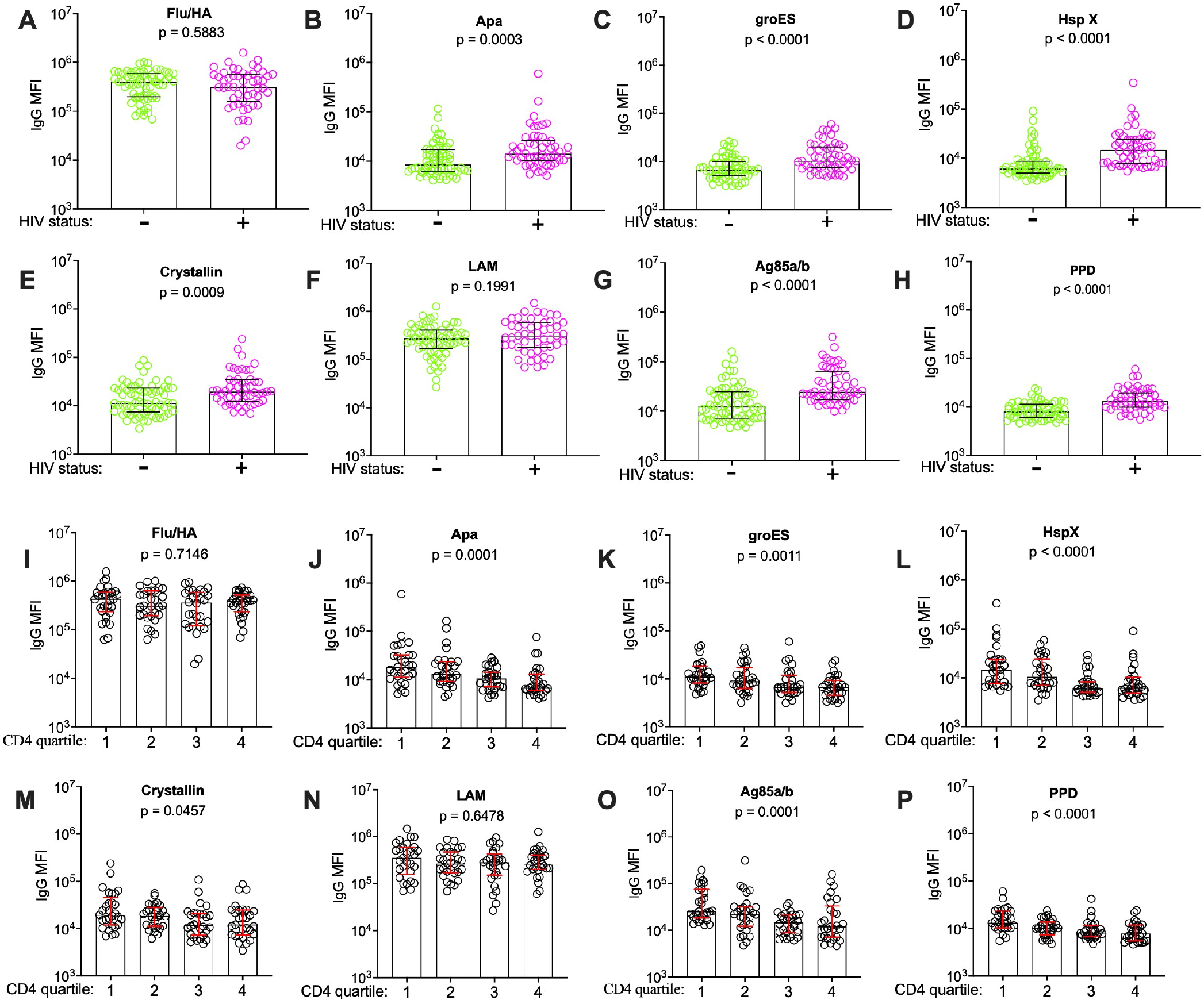
Significantly higher *Mtb*-related total IgG antibodies in the lowest CD4 T cell count quartile. **A-H** Influenza HA IgG and multiple *Mtb*-specific antigens IgG expression by HIV status (green: HIV− and pink: HIV+). **I-P** Influenza HA IgG and multiple *Mtb*-specific antigens IgG expression across CD4 count quartiles in all study participants and Mann-Whitney *U* and Kruskal-Wallis test for statistical significance were used, respectively. (n=115, HIV− n= 64, HIV+ n= 51)

### Non-specific hypergammaglobulinemia contributes to, but does not entirely account for the high anti-*Mtb* antibody levels in HIV-positive people with low CD4 cell counts

Hypergammaglobulinemia is a known consequence of HIV infection and is driven by non-specific activation of T cells and plasma cells (18, 19). To determine whether this accounted for the observed results, we used a quantitative ELISA to measure total IgG in all study participants. Total IgG levels were significantly higher in HIV-positive compared to HIV-negative participants (*p* = 0.0001) (Figure 4A), consistent with hypergammaglobulinemia in the HIV-positive group. We next correlated total IgG levels with *Mtb*-specific IgG antibody levels in all participants, and in participants grouped according to HIV status (Figure 4B). When evaluated across all participants, Spearman’s correlations of total IgG to specific anti*-Mtb* antigens IgGs showed inconsistent results for the *Mtb*-specific antigens. There was a significant positive correlation for some *Mtb*-related antigens i.e. HspX-specific IgG (*r* = 0.3337, *p* = 0.0003), Ag85a/b-specific IgG (*r* = 0.3101, *p* = 0.0009), PPD IgG-specific (*r* = 0.3216, *p* = 0.0006), ESAT-6/CFP-10-specific IgG (*r* = 0.2152, *p* = 0.0233), while others including Apa-specific IgG, GroES-specific IgG, Crystallin-specific IgG, LAM-specific IgG (and the non-*Mtb* control Flu/HA-specific IgG,) did not correlate with total IgG titer. (Figure 4B). The absence of correlation of total IgG with influenza HA-specific IgG, in contrast to positive correlations seen in multiple *Mtb* antigens, suggests an *Mtb*-specific process, that is not adequately explained by hypergammaglobulinemia alone. However, when participants were stratified by HIV status, there was no significant correlation between total IgG and any *Mtb*-specific antibodies, suggesting that those correlations observed in the unstratified group were driven by HIV status. Thus, HIV-mediated hypergammaglobulinemia does not appear to entirely explain the increased *Mtb*-specific IgG levels found in HIV-positive individuals in this cohort.

**Figure 4:**
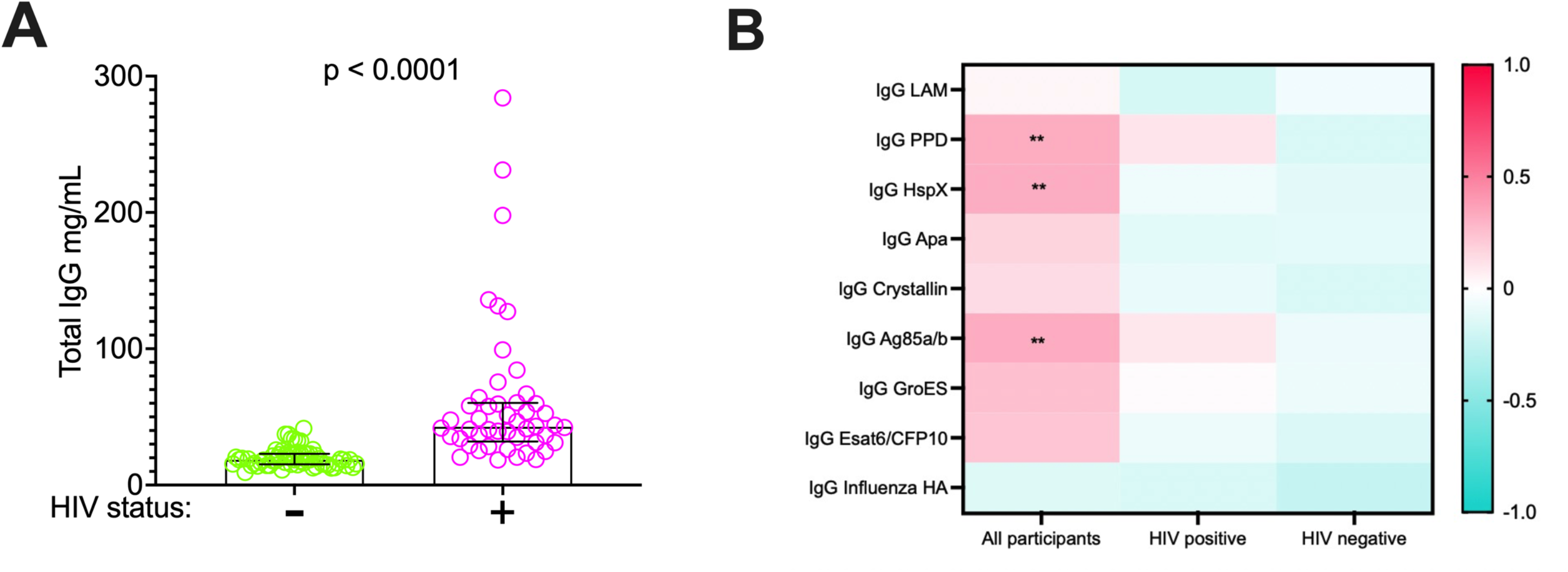
Total IgG positively correlate with most of the *Mtb*-related IgGs. **A** Total IgG content stratified by HIV status and Mann-Whitney U test used to test significance between the groups. **B** Heatmap showing multiple anti-*Mtb*-related IgG and total IgG correlations measured using Benjamini-Hochberg correction.

### Lowest ESAT-6/CFP-10 IFN-γ production but highest ESAT-6/CFP-10 IgG in HIV-positive people with low CD4 cell counts

The QuantiFERON assay measures *Mtb*-specific T cell immune response to specific *Mtb* antigens ESAT-6/CFP-10 by quantifying IFN-γ produced in response to these antigens. Measurement of ESAT-6/CFP-10-specific IFN-γ and IgG in the same individuals, stratified by CD4 cell count quartile demonstrated that people in the lowest CD4 count quartile produced the lowest levels of *Mtb*-specific IFN-γ (*p* = 0.0005) and the highest levels of *Mtb*-specific IgG (*p* = 0.0001) (Figure 5).

**Figure 5:**
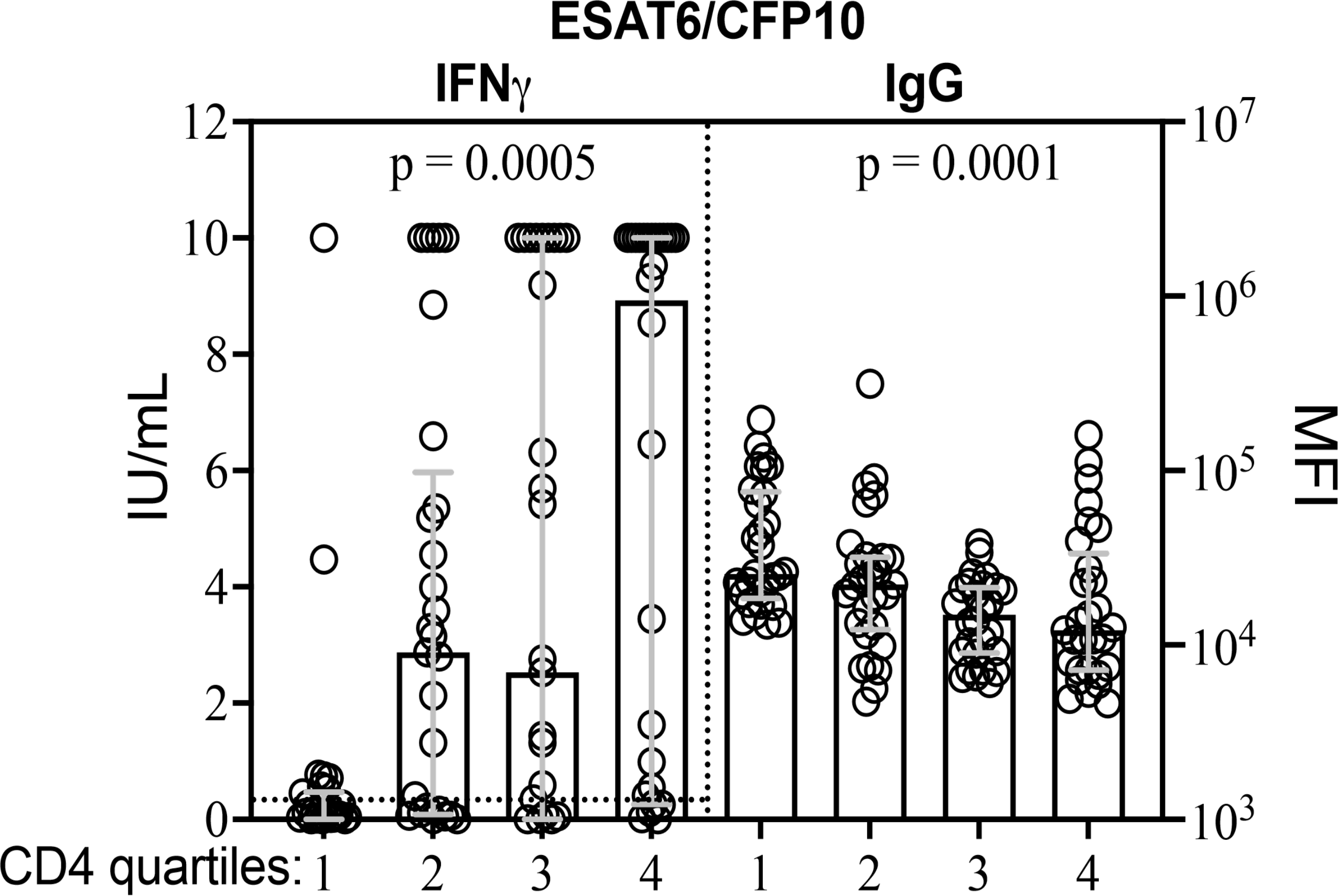
Discordant ESAT-6/CFP-10 specific IFN-γ versus ESAT-6/CFP-10 IgG expression across all participants CD4 count quartiles. Left panel: QuantiFERON (QFT) assay panel showing ESAT-6/CFP-10 specific IFN-γ across all CD4 count quartiles (1 = 0 - 428: 2 = 429 - 685: 3 = 686 - 923: 4 = 924 - 2043 in cells/μ L) (n = 118). Kruskal-Wallis test was used to determine statistical significance and the dotted line on this panel represents the positive test cut off (</= 0.35 IU/mL) for the QFT assay. Right panel: Luminex based antibody assay panel showing ESAT-6/CFP-10-specific total IgG antibodies across all CD4 count quartiles (1 = 0 - 428: 2 = 429 - 685: 3 = 686 - 923: 4 = 924 - 2043 all in cells/μ L) (n = 115). Kruskal-Wallis test was used to determine statistical significance.

## Discussion

In a group of well characterized of HIV-negative and antiretroviral-naïve HIV-positive people from KZN, South Africa we identified discordant results between levels of *Mtb*-specific IFN-γ and IgG, which was most striking in HIV-positive people with low CD4 cell counts. We found that 58% of adults in our study population were latently infected with *Mtb*, as defined by a positive QFT test. Male sex, older age and higher CD4 count were associated with QFT positivity. In contrast, in the same group of people, ESAT-6/CFP-10*-*specific IgG was significantly higher in people who were HIV-positive and had low CD4 cell counts. Notably, in participants in the lowest CD4 count quartile, levels of ESAT-6/CFP-10-specific IFN-γ released were lowest and levels of anti-ESAT-6/CFP-10*-*specific IgG antibodies were highest (Figure 5). Antibodies to 6 of 7 *Mtb*-specific antibodies followed this same pattern, while a control antibody specific to the hemagglutinin protein of influenza, did not. These results suggest that the QFT assay may underestimate LTBI among immunosuppressed people with HIV and *Mtb*-specific IgG may be a useful alternative biomarker for *Mtb* infection.

The current gold standard for the diagnosis of LTBI are interferon gamma release assays like the QFT, which rely on host T cell function to produce IFN-γ in response to *ex vivo* stimulation. Interestingly, in our study HIV status was not an independent modulator of QFT positivity, while CD4 count was. Across the entire study population, there appeared to be no difference between QFT rates in HIV-negative and HIV-positive participants. By stratifying participants into CD4 count quartiles, we observed that HIV-positive people with relatively preserved CD4 counts (429 – 923 cells/mm^3^) had higher QFT positivity when compared to HIV-negative individuals with similar CD4 counts. Quantitative analysis of IFN-γ production in response to ESAT-6/CFP-10 showed that HIV-positive participants in the lowest CD4 quartile produced significantly lower levels IFN-γ compared to HIV-positive people in the higher CD4 count quartiles. In the absence of plausible explanation behind why people with lower CD4 cell counts, would have less exposure to *Mtb* infection, it is likely that these results are due to CD4 T cell depletion and/ or exhaustion causing false-negative QFT tests in this group. Our QFT positivity results are in line with previous studies that have found lower rates of IGRA positivity in people with lower CD4 T cell count (12, 20) and the underestimation of QFT positivity in PLWH is consistent with prior literature (21, 22).

We also found that increasing age was an independent predictor of QFT positivity and that is consistent with a cumulative *Mtb* exposure over years of life (20, 23). Male sex was also an independent predictor of QFT positivity in this study and is consistent with reports from other settings (24, 25). The mechanisms for higher rates of *Mtb* infection and TB disease in males is not precisely defined, but is thought to be due to a combination of social factors that may promote *Mtb* exposure, such as environmental and occupational exposures and tobacco use, and increased biological susceptibility due to the differential effects of sex hormones on TB immunity (24–27).

IgG antibodies against *Mtb*-related antigens were considered potentially useful markers for *Mtb* infection over a decade ago, but further development of diagnostic modalities utilizing them was discouraged by WHO (2012) in favor of using the available methods at that time (28). Here, using a customized Luminex assay (17), we found that HIV-positive individuals had higher levels of ESAT-6/CFP-10 IgG compared to HIV-negative people; this finding contrasted with the similar QFT results observed in HIV-positive and HIV-negative groups. Older age and decreased CD4 count were also associated with increased ESAT-6/CFP-10-specific IgG, congruent with the QFT results. For QFT, we observed the lowest rates of positivity in individuals in the lowest CD4 count quartile, in contrast to the *Mtb*-specific plasma circulating anti-ESAT-6/CFP-10-specific IgG antibodies which had the highest concentration in this group (*p* = 0.0001) (Figure 5). Antibodies to 6 of the 7 *Mtb* antigens assessed followed the same pattern as anti-ESAT-6/CFP-10-specific IgGs while the non-*Mtb*-related IgG control (the hemagglutinin protein of influenza) did not vary between HIV or CD4 groups, suggesting that the finding of higher *Mtb*-specific antibodies was pathogen-specific. Interestingly, LAM-specific IgG was not influenced by HIV infection or low CD4 count. A possible reason for this is that LAM is not specific to *Mtb*, and is found in the cell wall of other species of *Mycobacterium* that are ubiquitous in the environment (29).

Hypergammaglobulinemia which has been linked to HIV infection (18) may be a non-specific reason for elevated Mtb-specific antibodies in PLWH. Consistent with prior literature, in this study, total IgG titers were higher in HIV-positive participants. Hypergammaglobulinemia may partially explain higher anti-*Mtb* antibodies in PLWH, but we observed contrasting patterns between hemagglutinin-specific (*Mtb* non-specific) and *Mtb*-specific IgG, indicating that hypergammaglobulinemia alone does not explain higher anti-*Mtb* antibody levels in those with low CD4 counts. Further experiments with independent and larger cohorts are required to determine whether elevated levels of anti-*Mtb*-related IgG can be detected independent of or when controlling for generalized hypergammaglobulinemia and whether detection of *Mtb*-specific antibodies may be a useful diagnostic strategy, particularly in HIV-positive participants with low CD4 counts. Overall, our results point to a potential diagnostic role for *Mtb*-specific antibodies to provide a way of defining *Mtb*-infection status that is not dependent on T cell function.

This study has several limitations. These data were collected from a convenience sample and are therefore not population representative, which limits generalizability. All HIV-positive individuals assessed were antiretroviral therapy naïve, which also limits generalizability. Limitations of the data include the lack of positive/negative cut-off thresholds or absolute quantification for the beads-based *Mtb*-specific IgG assays used here. The assay also measures relative concentration rather than an absolute quantitative measure of concentration that would allow direct comparison to total IgG concentrations, which would be required to address the contribution of hypergammaglobulinemia more definitively. Future work that combines well-defined HIV-positive (including people on antiretroviral therapy) and HIV-negative individuals from areas of high and low TB endemicity will be needed to advance the evaluation of *Mtb*-specific IgGs as alternative biomarkers for LTBI detection.

Even with its limitations and possible under-estimate of LTBI in HIV-positive people with low CD4 counts, our finding of QFT-confirmed LTBI in 58% of 18-50 years old adults is higher than has previously been reported in a generalized South African population. In a previous study using TST, only 34.3% of adults in South African urban townships around Johannesburg were reported to have LTBI (23). Mathematical models estimation using TST data supports the above results, reporting 30-40% South Africans having LTBI according to those estimations (30). A TST survey among gold miners in Johannesburg showed LTBI rates as high as 85%, which correlates with the higher levels of active TB cases identified within gold miners (31). A survey in a Cape Town township showed that about 40-50% of adolescents had LTBI using QFT assay (32), in line with the rates we report here. However, a survey of adolescents in rural KZN using QFT showed a much lower rate of 23% LTBI (33). With high rates of latent TB infection, South Africa is prioritizing preventative therapies, and to this end less expensive, easier and more accurate tests for LTBI especially in HIV-positive people are urgently needed (34).

Overall, our data suggest high rates of latent TB infection among HIV-negative and HIV-positive adults and that the QFT assay likely underestimates LTBI in HIV-positive people with low CD4 count. Eliminating TB remains challenging due to the limitations in diagnosing TB at various stages along the infection spectrum (2). Furthermore, while increasing access to TB preventative therapy is a global priority, very limited screening for LTBI in South Africa has been done, in part because of the indirect nature and complexity of interpretation required by the current assays. *Mtb*-specific IgG antibodies may hold promise as alternative biomarkers for LTBI, especially in people with low CD4 count.

## Acknowledgments

We appreciatively acknowledge the following: the Phefumula study participants who were all based at the KwaDabeka Clinic in Durban, KwaZulu-Natal, South Africa; the Africa Health Research Institute (AHRI) Clinical core study team for recruiting participants and collecting samples in KwaDabeka clinic; AHRI Biorepository team for their assistance with sample transport to AHRI laboratories, sample distribution and long-term storage of samples and the AHRI Data team for facilitating data collection and data management.

## Data sharing

Deidentified participants data used to generate results for this work will be available upon request. Requests for data can be sent to

## Contributors

MM, TN and EBW designed the overall study. ZM, DR and FK collected the clinical samples. MM, TB and SK performed the QuantiFERON assays. KAB and LRLD performed the antibody assays. MM, EBW, KB, LRLD, GA and TN performed the data analyses and interpretation, and LM validated all the statistical methods used. EBW, TN and GA provided supervision. MM, TN, and EBW wrote the manuscript. All authors reviewed the manuscript and approved the final version.

